# Microtubule binding protein Togaram1 is required for proper development of mammalian forebrain and neural primary cilia

**DOI:** 10.64898/2026.04.13.717734

**Authors:** Clarissa Q. Nassar, Savera J. Shetty, Noelle D. Dwyer

## Abstract

Proper forebrain development relies on precise spatial and temporal control of early neural stem cell (NSC) proliferation and later neurogenesis. Brain malformations can arise when these processes are defective. Joubert Syndrome (JS) is a neurodevelopmental disorder that is diagnosed by a mid-hindbrain malformation, but often includes forebrain defects such as microcephaly, which are less understood. One gene recently linked to Joubert Syndrome with microcephaly is *Togaram1*, which encodes a TOG domain microtubule binding protein shown to affect primary cilia. In the embryonic dorsal forebrain, NSCs have primary cilia on their apical membranes that play a role in regulating proliferation and neurogenesis, but how they do this is not well understood. Here we investigate the role of *Togaram1* in mammalian forebrain development using a mouse knockout. We find that *Togaram1* is crucial for forebrain size, thickness, and morphology. In particular, knockout forebrains have sporadic indentations of the lateral ventricles, and the neuronal layer is thin with gaps and heterotopias. The dorsal forebrain NSCs have increased proliferation and apoptosis. Finally, the primary cilia of *Togaram1* knockout NSCs have abnormal morphology and function. This study begins to elucidate the role of *Togaram1* in forebrain morphogenesis and the involvement of NSC primary cilia in forebrain malformations.

## INTRODUCTION

The mammalian forebrain develops from the most rostral end of the neural tube to form some of the most complex parts of the brain such as cerebral cortex, basal ganglia, hippocampus, and thalamus. It starts as a pseudostratified epithelium made up of neural stem cells (NSCs). NSCs are elongated polarized epithelial cells that have basal endfeet near the pia and apical endfeet at the ventricle, joined by adherens junctions. Early in development these cells proliferate exponentially to increase the area of the neuroepithelium and form the cerebral hemispheres. Later, they divide asymmetrically to produce neuronal daughter cells that migrate away from the ventricle and do not divide again. Disruptions to NSC proliferation or neurogenesis can cause brain malformations and neurodevelopmental abnormalities (McNeely & Dwyer, 2021; Thomsen et al., 2025; Suciu & Caspary, 2021). How these processes are precisely regulated during healthy brain development is a topic of intense study.

The NSCs each have a non-motile primary cilium on their apical membranes that is thought to sense developmental signals. Shh signaling mediated by primary cilia is important for both neural patterning and proliferation in the nervous system (Cai et al., 2023). Cilia-related mutations cause a class of disorders known as ciliopathies that can affect various organs in the body such as limbs, eyes, and brain. Joubert Syndrome (JS) is a neurodevelopmental ciliopathy that is diagnosed based on a mid-hindbrain malformation termed the molar tooth sign, but other brain malformations including microcephaly can occur (Kumandas et al., 2004; Latour et al., 2020; Morbidoni et al., 2021).

*TOGARAM1* (TOG array regulator of axonemal microtubules 1) is one of forty genes that causes JS, but its function is not well understood. Mutations in it have been found in several families with affected children or fetuses; microcephaly was observed in some, and cognitive disability in others occur (Latour et al., 2020; Morbidoni et al., 2021). The Togaram1 protein (aka FAM179B or Crescerin1) contains 4 tumor overexpression gene (TOG) domains, which have been shown to bind and polymerize microtubules (Das et al., 2015). In in vitro microtubule polymerization assays, Togaram1 protein supported slow microtubule growth (Das et al., 2015; Saunders et al., 2025). Loss of Togaram1 in zebrafish results in short cilia with decreased post-translational modifications and body defects, such as scoliosis and kidney cysts (Latour et al., 2020). Additionally, structural and functional cilia defects are also observed in mutants of Togaram1 homologs in C. elegans, Chlamydomonas, and Tetrahymena (Louka et al., 2018; Morbidoni et al., 2021; Perlaza et al., 2022).

Here we use a mouse knockout model to better understand the role of *Togaram1* in forebrain development and Joubert Syndrome. Although *Togaram1* mutations in the mouse were previously shown to cause lethality, microphthalmia, and neural tube defects, its requirement in forebrain development was not analyzed (Chee et al., 2023; Wang et al., 2025). Here we find that *Togaram1* knockout embryos at the gross level have smaller rounder cerebral hemispheres, microphthalmia, and polydactyly, which are hallmarks of JS and other ciliopathies. Analyses of forebrain development through sectioning and immunohistochemistry show morphological abnormalities at the lateral ventricle surface and the nascent neuron layer. In dorsal forebrain, we observe a thinner cortex, abnormal neurogenesis, displaced mitotic cells, and increased apoptosis. Finally, we find that primary cilia of NSCs in the knockout forebrains are malformed and have abnormal processing of Shh signaling component Gli3. This study advances our understanding of *Togaram1* requirement in forebrain development and provides new insight into the pathogenesis of the microcephaly in Joubert Syndrome.

## RESULTS

### Togaram1 loss reduces embryonic survival and causes developmental abnormalities

To ascertain the requirements for Togaram1 in mammalian development, we obtained a *Togaram1* knockout mouse line from The Jackson Laboratory. This line was generated using Crispr-Cas9 to delete 542 bp including the second coding exon of the *Togaram1* gene (**Fig. 1A**). It is predicted to be a null allele by causing a frameshift after 684 amino acids followed by premature stop codons, leading to either nonsense-mediated decay, or at best an N-terminal fragment. We confirmed the gene deletion using whole genomic sequencing. Published data from single-cell RNAseq databases suggests that *Togaram1* mRNA is expressed in the nervous system structures of early mouse embryos, including cortical NSCs, intermediate progenitors, and immature neurons (Di Bella et al., 2021; La Manno et al., 2021). To test for loss of *Togaram1* mRNA in the knockout, we used quantitative RT-PCR and found that *Togaram1* mRNA is reduced in heterozygotes and undetectable in homozygous knockout tissues, but is present in control brains as well as liver (**Fig. 1B**). Immunoblot testing for Togaram1 protein was not possible because the eight different antibodies tested either did not detect any protein or they showed non-specific bands (see Methods).

**Figure 1.**
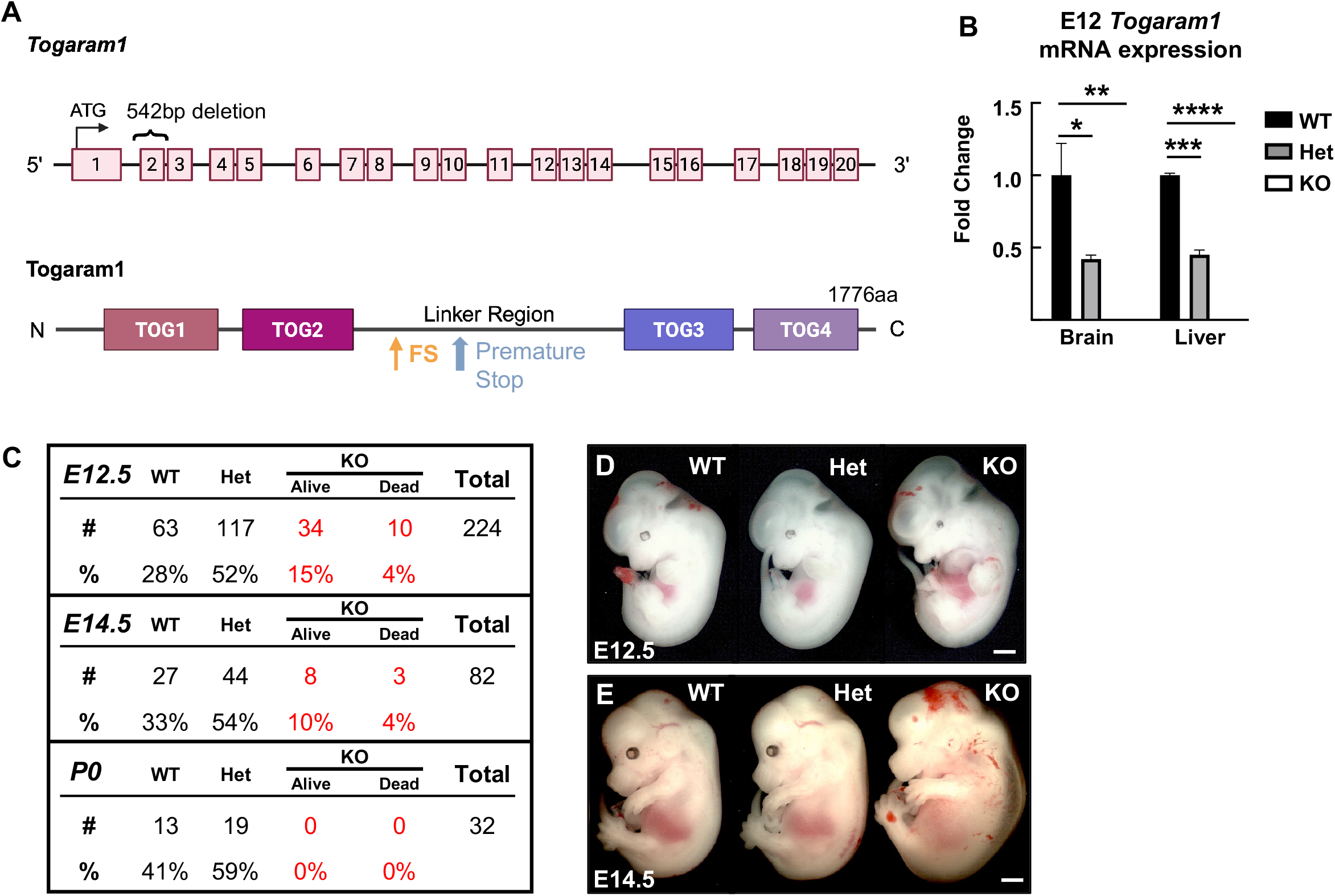
*Togaram1* mouse KO reduces embryonic survival and disrupts proper craniofacial development. (A)Schematic of *Togaram1* gene and protein. (B) *Togaram1* mRNA is undetectable in knockout (KO) embryos and reduced in heterozygous (Het) embryos at embryonic day 12.5 (E12.5) brain and liver tissues using q-RT-PCR. Fold change values are normalized to WT. n = 3 embryos per genotype for each tissue. One way ANOVA. (C) Progeny of Het x Het crosses collected at E12.5, E14.5, and P0. E12.5 and E14.5 KO embryos found at reduced Mendelian ratio. (D, E) Example images of Togaram1 WT, Het, and KO embryos at age E12.5 and E14.5. Scale bar = 1mm.**15/15**

Next, we assessed the survival of homozygous knockout animals by collecting progeny from heterozygous x heterozygous crosses. Heterozygotes are viable and fertile, and appear normal. We were not able to collect any homozygous knockouts at postnatal day 0 (P0), indicating prenatal lethality. To determine the age of lethality, we collected embryos from heterozygous crosses. At embryonic days (E) 12.5 and 14.5, live knockout embryos can be found, but at reduced Mendelian ratios of 15% and 10%, respectively, rather than the expected 25% (**Fig. 1C**). The *Togaram1* knockout embryos found alive at E12.5 and E14.5 display gross head and limb abnormalities, but have no significant difference in body size; meanwhile heterozygous embryos are indistinguishable from wild-types (**Fig. 1D, E**).

The *Togaram1* homozygous knockouts have grossly visible phenotypes of the brain, eye, face, heart, and limbs. All knockout embryos collected have microphthalmia and small malformed forebrains (**Fig. 2A**). Microcephaly is seen in 70% of knockout embryos and the remaining 30% have exencephaly (**Fig. 2A, C**). Additionally, roughly 50% of knockout embryos display flipped heart orientation, indicative of laterality defects (**Fig. 2B**). *Togaram1* knockouts also have cleft lip and rear paw preaxial polydactyly visible at E14.5 (**Fig. 2D, E**). These data indicate that *Togaram1* loss reduces embryonic viability in late gestation, abolishes postnatal survival, and disrupts development of the brain and other organs. For the remainder of this paper, we focus on the forebrain at age E12.5, the latest age when live embryos could be readily collected in sufficient numbers for quantitative analyses.

**Figure 2.**
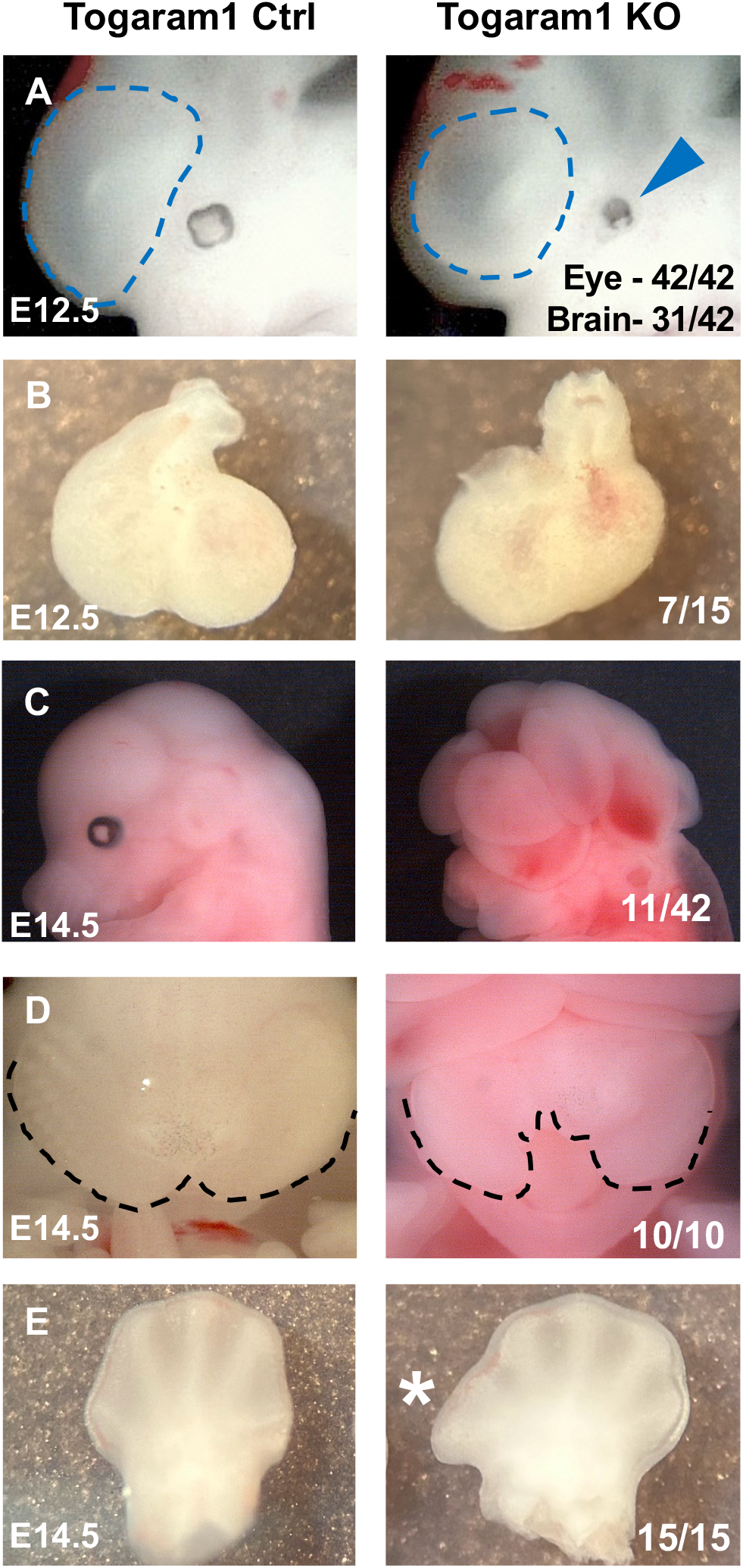
Brain, craniofacial, heart, and digit abnormalities found in *Togaram1* KO. (A-E) Representative images of phenotypes found in *Togaram1* KO at ages E12.5 and E14.5. Fractions on right are KO embryos with each phenotype out of total (combined E12.5 and E14.5). (A) Microcephaly (forebrain outlined by dashed blue line) and microphthalmia (blue arrowhead) are observed in KO. (B)Heart laterality is randomized in KO. (C)Exencephaly is observed in ∼ ¼ of KO brains. (D)KO have cleft lip (marked by dashed line). (E)Pre-axial polydactyly on rear paws at E14.5. Star labels extra digit.GE

### Togaram1 is required for normal forebrain size and morphology

To determine the role of Togaram1 in forebrain development, we analyzed cerebral hemisphere size and morphology. As we had previously observed a small forebrain within the skull, we dissected out E12.5 brains to measure their area, length, and width. The knockouts have a reduced hemisphere area and length, but no change to the hemisphere width (**Fig. 3A, B**). Furthermore, the knockout brains appear more transparent (**Fig. 3A**), and indeed measuring cortical thickness on cross sections showed the knockout cortices are significantly thinner (**Fig. 3C)**. To further investigate forebrain morphology, we compared coronal forebrain sections spanning rostral to caudal planes from controls and *Togaram1* knockouts. Control brains have a smooth dorsal forebrain (nascent cortex), and two ganglionic eminences (MGE, LGE) in the ventral forebrain (nascent basal ganglia) (**Fig. 3D, ctrl**). While *Togaram1* knockout brains appear to have overall proper dorso-ventral organization, they have abnormal ventricle shapes with variable numbers of indentations of the ventricular surface (**Fig. 3D, KO, dashed boxes**). These ventricle indents were found on 13 out of 18 dorsal hemispheres analyzed, and their depth was highly variable, ranging from 35um to 156um (**Fig. 3D-dorsal, 3E, F**). The ventral forebrain of knockouts had an altered number and structure of ganglionic eminences in 17 out of 18 ventral hemispheres examined, with up to 5 eminences present in one section (**Fig. 3D-ventral, 3G**). These data demonstrate that Togaram1 is crucial for proper forebrain size and structure.

**Figure 3.**
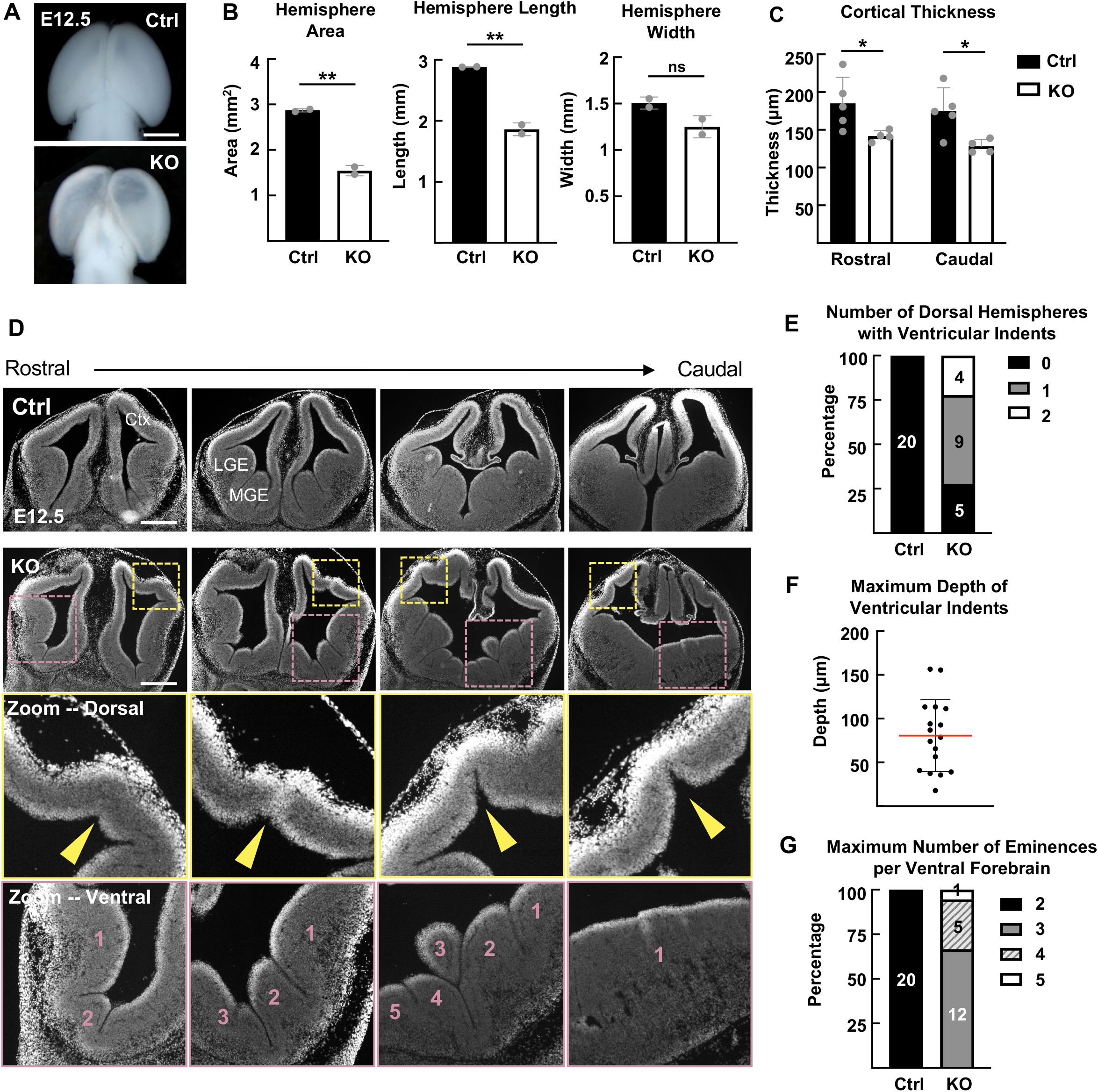
Forebrain architecture is altered in *Togaram1* KO. (A) Representative dorsal views of E12.5 control and KO forebrains. (B) KO forebrain shows reduced average area and length, while width is not significantly affected at E12.5. (C) Quantification of average cortical thickness shows significant difference between control and KO embryos in rostral and caudal sections. (D)Serial coronal forebrain sections spanning from rostral to caudal regions reveal altered morphology in E12.5 KO. Zoom of dorsal region shows the ventricular indentations of the cortex. Zoom of ventral region shows the presence of extra ganglionic eminences. Ctx, Cortex; MGE, medial ganglionic eminence; LGE, lateral ganglionic eminence. (E) KO forebrain hemispheres have variable numbers of ventricular indents. (F) Maximum depth of ventricular indents found in KO. (G) KO forebrain hemispheres have variable extra ganglionic eminences observed per hemisphere in KOs. For B, n = 2 brains per genotype; unpaired T-test. For C, n = 5 control, 4 KO brains; unpaired T-test. For E-G, n = 20 control and 18 KO hemispheres. Numbers inside graphs indicate number of hemispheres in that category. Scale bars: 1 cm in A; 500 µm in D.

### Togaram1 deletion disrupts neuronal layer formation in early cerebral cortex

Given the findings of ventricular indentations and thinner cortex, we wondered whether the neuronal layer forms properly in the *Togaram1* knockout brains. We stained coronal sections for the neuronal tubulin Tubb3 to visualize the developing neuronal layer in E12.5 brains (called the preplate at this stage). In control cortices, a thin but continuous Tubb3+ cell layer overlies the ventricular zone of NSC nuclei (**Fig. 4A**). However, in the knockout brains, the neuronal layer and the phenotype was variable. In some KO brains, a continuous neuronal layer was present, but thinner above a ventricular indentation (**Fig. 4B**). In other KO cortices, the neuron layer was very thin and discontinuous with gaps (**Fig. 4C**). Furthermore, abnormal clumps of neurons (neuronal heterotopias) of variable size and position were sporadically seen in the dorsal forebrains of *Togaram1* knockouts (**Fig. 4B, 4C**). These findings indicate that Togaram1 is required for normal cortical neurogenesis including sufficient numbers and layering, but is not required for cortical neuron radial migration away from their birthplace at the ventricular surface.

**Figure 4.**
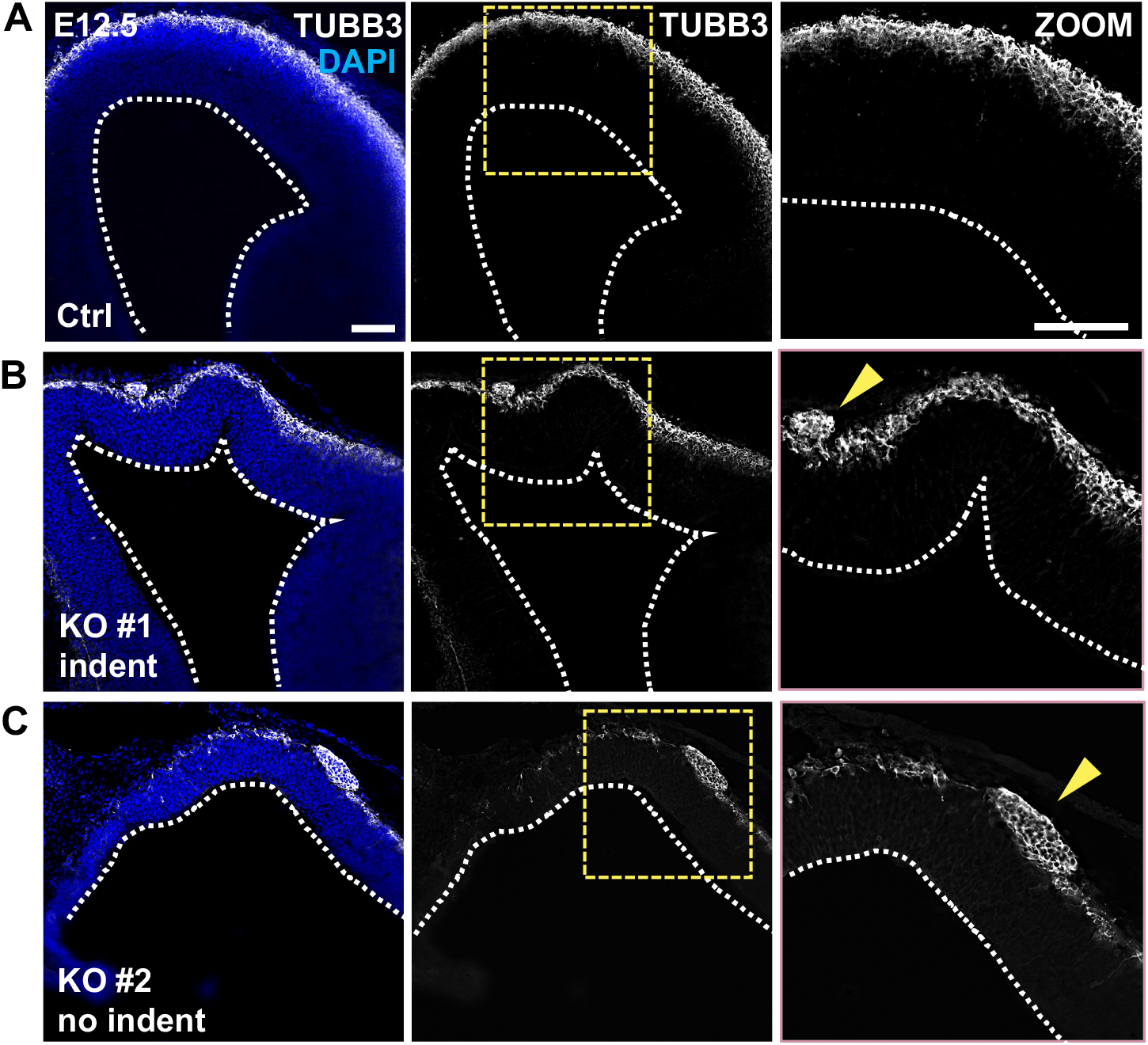
Neuronal layer is thinner and irregular in *Togaram1* KO dorsal forebrain. (A)Control cerebral cortex section shows a continuous neuronal layer of even thickness at E12.5, seen stained with DAPI and neuronal tubulin (Tubb3). (B, C) Two different KO forebrains show variable defects of the neuron layer. (B)KO #1 cortex has a ventricular indentation, thinning of the neuron layer above the indent, and abnormal neuronal cluster (arrowhead). (C)KO #2 cortex does not have ventricle indent but a very thin neuron layer with multiple gaps and an abnormal cluster (arrowhead). White dashed line marks ventricle. Scale bar = 100um.

We hypothesized that the ventricular indentations and thin neuron layer of the knockout brains could result from excess NSC mitoses at the ventricular surface, and/or excess apoptosis. To test whether mitotic index is increased or the position of mitotic cells is altered in the knockout brains, we stained for phospho-histone H3 (PH3). The overall mitotic index is significantly increased in the knockout cortices, mostly due to an increase in mitotic nuclei away from the ventricle (abventricular) (**Fig. 5A, B**). To compare apoptosis levels in control and *Togaram1* knockout brains, we immunostaned for cleaved caspase 3 (CC3). While apoptosis is very low in wild-type cortices at E12.5, it is increased fourfold in the knockout brains. The apoptotic cells are found in the neuronal layer as well as the stem cell layer (**Fig. 5C, D**). These data suggest that loss of Togaram1 affects mitotic duration or positioning of nuclei during mitosis, and causes an increase in apoptosis of both NSCs and neurons.

**Figure 5.**
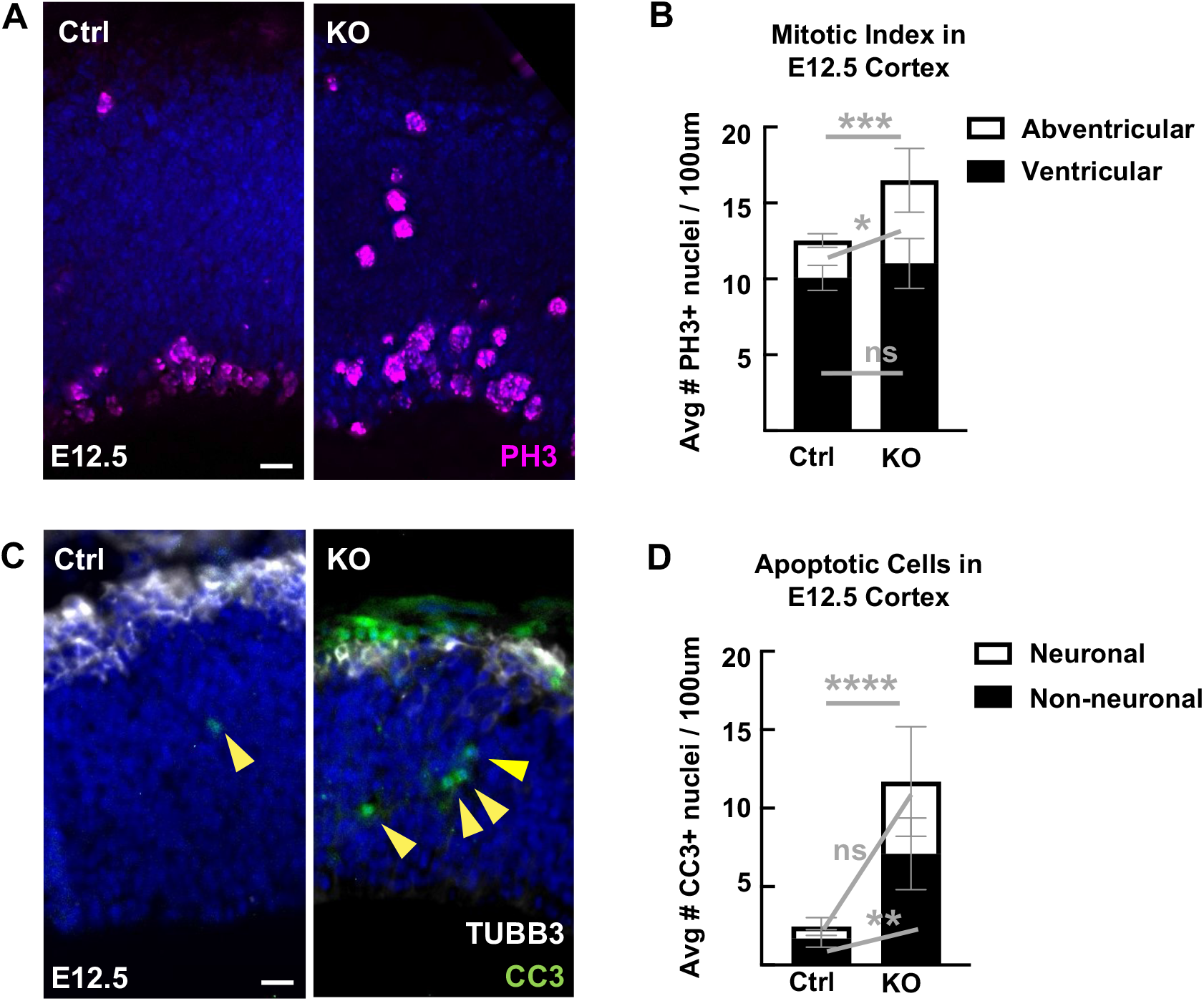
Mitotic index and apoptosis is altered in Togaram1 KO. (A)Control and KO E12.5 cortical section stained with DAPI (blue) and phospho-Histone H3 (PH3, magenta). (B)Quantifications of PH3+ cells in control and KO sections shows an increase in the number of PH3+ cells away from the ventricle. (C-D) Apoptosis is increased in non-neuronal and neuronal cells of E12.5 KO cortices stained for DAPI (blue), cleaved Caspase-3 (CC3 green, arrowheads), and Tubb3 (white). Scale bars = 20um. For B, D, n = 4 control and 4 KO brains (4 sections averaged for each); Unpaired T-test.

### Togaram1 alters cilia formation and function in the developing forebrain

Since many Joubert Syndrome genes are thought to affect primary cilia, and *Togaram1* loss was shown to affect cilia in other model organisms (Das et al., 2015; Latour et al., 2020; Louka et al., 2018; Morbidoni et al., 2021; Perlaza et al., 2022), we hypothesized that *Togaram1* loss could affect the primary cilia of the forebrain. To image the apical endfeet of dorsal forebrain NSCs and their primary cilia *en face*, we prepared cortical slabs. We immunostained with antibodies against ZO-1 to label apical junctions and Arl13b to mark primary cilia. We noticed that the control brain NSCs had mostly linear cilia with a number of round cilia, whereas the knockout brain NSCs had mostly round cilia with only a few appearing linear (**Fig. 6A**). In addition to their abnormal shape, the length of the knockout cilia was significantly shorter (**Fig. 6B**). Next, since the cortical neuron layer is thin and irregular in the knockouts, we wanted to know if the primary cilia on neurons are also affected. We stained forebrain cross sections for ciliary marker Arl13b and centrosome marker gamma tubulin to visualize cilia in the neuronal layer. Indeed, there is a significant reduction in the length of primary cilia in the neuron layer although less severe than the NSCs (**Fig. 6C, 6D**).

**Figure 6.**
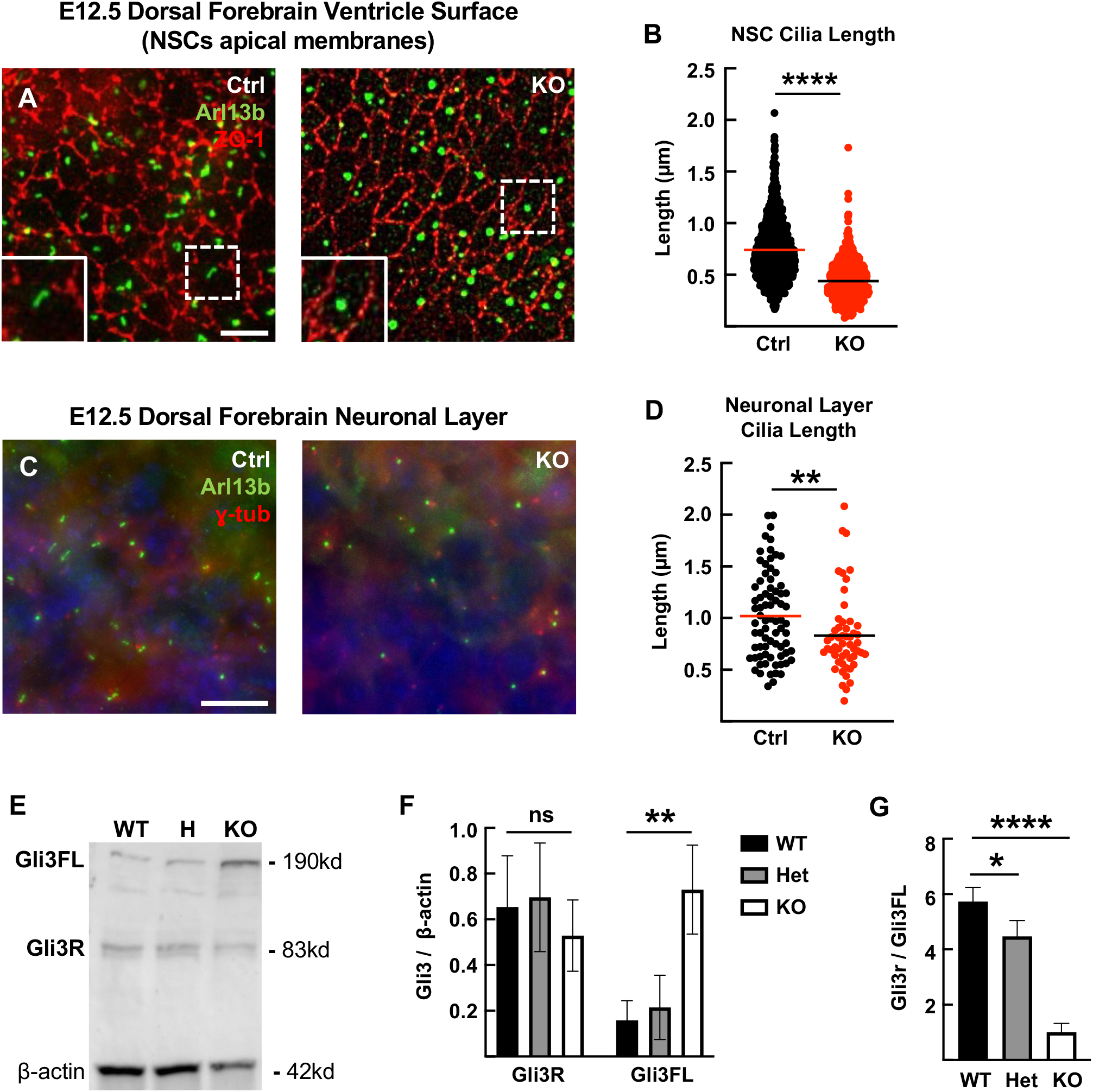
*Togaram1* KO primary cilia in dorsal forebrain have abnormal morphology and SHH signaling activity. (A)Representative images of dorsal forebrain ventricular surfaces showing NSCs apical membranes stained for Arl13B (green, cilia) and ZO-1 (red, apical junctions). Arl13b staining shows a linear shape in mouse control NSCs but a round shape in most Togaram1 KO NSCs. (B)NSC cilia length shows significant difference between control and KO groups. (C)Representative images of dorsal forebrain neuronal layer stained for Arl13B (green, cilia) and gamma tubulin (red, centrosomes). (D)Control and knockout neuronal cilia shows significant difference in length. (E)Western blot showing band intensities of full-length Gli3 (Gli3FL), repressor form Gli3 (Gli3R), and β-actin in whole brains collected from embryonic day 12.5 (E12.5) wild-type (WT), heterozygous (H), and Togaram1 knockout (KO) embryos. (F) Quantification of Gli3FL or Gli3R levels normalized to β-actin shows increase in full-length Gli3 in KO brains. Bars represent mean ± s.e.m. (G) Ratio of Gli3R to Gli3FL is significantly decreased in KO brains compared to controls. For B, n = 1032 control and 1043 knockout cilia. For D, n = 76 control and 53 knockout cilia. For B and D, Kolmogorov-Smirnov (K.S.) and Mann-Whitney (M.W.) tests. For E-G, n = 3 brains per genotype. One way ANOVA. Scale bars: 10 µm in A and D.

To test for disruption of the function of primary cilia in *Togaram1* knockout brains, Sonic hedgehog (Shh) signaling was assessed. Shh signaling is involved in forebrain growth and patterning and is the best understood signaling pathway mediated by embryonic primary cilia. Gli3 is one of its main downstream effectors and the cleavage of full-length (FL) Gli3 into Gli3 repressor (Gli3R) plays a key role in the development of the dorsal forebrain. Therefore, we immunoblotted for the Gli3 protein in control and knockout brain lysates (**Fig. 6E-G**). Knockout brains have an increase in the amount of Gli3FL, and a decrease in the Gli3R/FL ratio. This disruption of normal Gli3 processing in the brain shows that *Togaram1* loss affects the function of NSC primary cilia as well as their morphology.

## DISCUSSION

How neural stem cells create and grow the brain is a fundamental question in developmental biology. Genes mutated in human brain malformations can provide valuable insights into this question. While ciliopathy syndromes often include forebrain malformations, the functions of primary cilia in forebrain development remains less well understood than in the hindbrain or spinal cord. Here we focus on the role of the Togaram1 gene in forebrain morphogenesis. While *Togaram1* was first identified as a causative gene for Joubert Syndrome, classically considered a mid-hindbrain malformation, we show here that Togaram1 is important for forebrain size, morphology, and neurogenesis. We find that in the mouse *Togaram1* knockout, the forebrain is present and has roughly normal shape, but the lateral ventricles have sporadic indentations of the ventricular surface. The neuronal layer is properly located superficial to the NSC layer, but is thin and discontinous, with occasional heterotopia. At the cellular level, we find excess apoptosis, abventricular mitoses, and malformed primary cilia in forebrain NSCs. The defective cilia are likely to cause the abnormal Hedgehog/Gli3 signaling, but how this could lead to misplaced mitoses or increased apoptosis is unclear. Much future work will be needed to determine whether all the morphological defects are secondary to the abnormal cilia, or whether Togaram1 affects multiple cellular processes such as cell division or migration.

Even within primary cilia, the precise function of Togaram1 is yet to be determined, with only a handful of publications so far in the literature. Detection of endogenous protein localization has been difficult, and here our efforts with eight commercial antibodies were fruitless. When epitope-tagged Togaram1 was overexpressed in HEK cells by another group, it showed localization on cytoplasmic microtubules as well as cilia (Das et al., 2015). However, the handful of publications with experimental data about Togaram1’s cellular function focused on either cilia morphology or purified microtubules in vitro. In in vitro microtubule assays, purified Togaram1 protein supported slow microtubule growth (Das et al., 2015; Saunders et al., 2025). It was suggested that TOG1 and 2 promote polymerization while TOGs 3 and 4 promote lattice binding (Das et al., 2015). Furthermore it was shown biochemically that Togaram1 binds in a multiprotein complex including Armc9 (Latour et al., 2020; Saunders et al., 2025). Cilia phenotypes in mutant model systems and cell lines were generally shorter, but varied with cell type and the specific mutation (Latour et al., 2020; Morbidoni et al., 2021; Perlaza et al., 2022; Wang et al., 2025) (Das et al., 2015; Louka et al., 2018). Some documented changes in tubulin postranslational modifications in mutant cilia (Latour et al., 2020), but the mechanism is unclear. Thus much more work in various systems and cell types is needed to understand the activities and interactions of this protein.

The mouse *Togaram1* knockout phenotypes we identified herein overlap with those found in humans with *TOGARAM1* mutations, but the human phenotypes varied. In one pedigree, two 2nd trimester fetuses with compound heterozygous mutations had microcephaly, cleft lip, and micropthalmia, but no polydactyly was noted (Morbidoni et al., 2021). In another study, four children with various *TOGARAM1* mutations had absence of cerebellar vermis, cognitive delays, and craniofacial defects, but not cleft lip or polydactyly, and forebrain size was not described (Latour et al., 2020). A third study identified a patient with spina bifida, intellectual disability, seizures, partial corpus callosum agenesis, and facial dysmorphism, but no cleft lip or polydactyly had a dominant missense mutation in *TOGRARAM1* (Wang et al., 2025). Thus it seems that the forebrain is usually affected when *Togaram1* is mutated, with phenotypes ranging from intellectual disability to structural defects detected if brain imaging was done. Our study provides a mouse model for the human forebrain defects, and begins to provide clues to the cellular etiology of these defects.

## MATERIALS AND METHODS

### Mice

Mouse colonies were maintained in accordance to the guidelines set by the National Institutes of Health and the policies approved by the Institutional Animals Care and Use Committee at the University of Virginia.

The mouse strain used for this research project, C57BL/6NJ-Togaram1em1(IMPC)J/Mmjax, RRID:MMRRC_046132-JAX, was obtained from the Mutant Mouse Resource and Research Center (MMRRC) at The Jackson Laboratory, an NIH-funded strain repository, and was donated to the MMRRC by Stephen Murray, Ph.D., The Jackson Laboratory (Adams et al., 2024). After obtaining this line, we backcrossed it 4 times and maintained it on a C57BL/6NCrl (Charles River) inbred background for breeding and experimentation. We verified the Togaram1 deletion using DNA sequencing and RT-PCR (see below).

Timed matings were set up, and the morning of the vaginal plug was considered E0.5. Littermate embryos were considered controls for all experiments. Mouse parent and embryo genotypes were confirmed via PCR. The sex of the embryos was not considered to be a biological variable relevant to our studies and therefore, it was not recorded. Embryos aged day 12 and day 14 were collected and the specific age of embryos used for each experiment is stated in the figure and/or its legend.

### Real Time PCR

Whole brains and liver tissue for the 3 genotypes were dissected from E12.5 embryos and snap frozen on dry ice. RNA was isolated using RNeasy Plus mini kit (Qiagen, cat #74134) according to the manufacturer’s handbook and quantified on a Nanodrop 2000. cDNA synthesis was done for 1ug of RNA/sample using Quantitect RT kit (Qiagen cat #205311). cDNA samples were diluted 15 times and real time PCR was performed for 1ul of diluted sample with PowerUp™ SYBR™ Green Master Mix (cat #A25742) using the delta Ct method on the ABI Step One Plus Instrument. A control reaction without reverse transcriptase was also set up to assess gDNA contamination.

Primers used:

Togaram1: F: 5’ AGA CCT CCA ATT CCA AGG ACA 3’

R: 5’ ATT ACC AGC ACC ACA TGT AGG AA 3’

Beta Actin: F: 5’ GAT GACCCA GAT CAT GTT TGA GAC C 3’

R: 5’ TAG TCT GTC AGG TCC CGG C 3’

### Western Blot

Whole brains from wild type, heterozygous and mutant Togaram1 embryos aged E12.5 were collected and snap frozen on dry ice. Lysates were prepared in RIPA buffer (pH7.4) with Halt™ Protease Inhibitor Cocktail (Thermo Scientific, cat #PI87786) using a manual pestle (Fisher Scientific, cat #12-141-364) and passing the homogenate through a 21g needle five times. After incubation on ice for 10 minutes the lysates were centrifuged at 14K RPM for 10 minutes at 40C. 20 micrograms of total protein, estimated by Pierce BCA Protein Assay Kit (ThermoFisher Scientific, cat #23227), were separated on 4-20% gradient gels and transferred to 0.45um nitrocellulose membrane (Thermofisher, cat #88018) at 100v for 90 minutes at 40C. Membrane was blocked in 5% milk TBS-T and incubated with goat anti-Gli3 (R&D Systems, cat #AF3690) at 1:1000 and mouse anti Beta Actin (SantaCruzTechnologies cat #Sc-47778) at a dilution of 1:5,000 in 5% milk TBS-T. Secondary antibodies were anti-goat HRP conjugated donkey IgG (R&D System, cat #HAF109) and anti-mouse HRP conjugated goat IgG (Millipore Sigma, cat #AP308P) at 1:10,000 dilution in 5% milk in TBS-T. The blot was developed with Pierce ECL substrate (cat #32209) and imaged on the Azure c300 imaging unit. Quantification of the bands were done with ImageJ.

### Cryosection preparation and immunostaining

Embryos were collected and their heads were pinched off and placed in 2% PFA for 4 to 6 hours at 4ºC. Heads were quickly washed in PBS and then moved to 30% sucrose at 4ºC until the heads sank. Heads were then embedded in OCT (Tissue-Tek, cat #4583) and stored at -80ºC until use. Sections were cut at 20µm on a Leica CM3050S cryostat and collected on Superfrost Plus slides (Fisher Scientific, cat #12–550-15). Cryosections were left at room temperature for 30-45 minutes to warm up. Slides were quickly rinsed with 1ml of PBS and then blocked with 2% Normal Goat Serum (Jackson Immuno, cat #AB-2336990) in PBST for one hour at room temperature. Sections were incubated with primary antibodies overnight at 4ºC. Sections were washed with PBS three times on the following day. Slides were incubated with secondary antibodies (1:200) and DAPI (4’,6-diamidino-2-phenylindole, 1:100) for 30 minutes at room temperature. Sections were washed quickly with PBS twice and then coverslipped with VectaShield fluorescent mounting medium (Vector Laboratories Inc., cat #H-1000).

### Cortical slab dissection and immunostaining

Slabs were prepared according to the protocol described in Janisch & Dwyer 2016. Embryos were decapitated and their skulls were opened to expose the forebrain. Heads were immersed in 2% PFA for 20 minutes. After this initial fixation, the cortices were pinched off and placed in a well with their apical side facing up. Slabs were trimmed until it was a flat surface and another 2% PFA fixation was done for 2 minutes. Next, slabs were incubated in PBS + 0.01% Triton (PBST) for 10 minutes before blocking for one hour with 5% Normal Goat Serum in PBST. Primary antibodies (see specifics below) were added and incubated for one hour at room temperature before being moved to 4ºC overnight. On the following day, slabs were washed once with PBST once and then twice washed with PBS. Secondary antibodies (1:200) and DAPI (1:100) were added to slabs and incubated for one hour at room temperature. Slabs were then washed twice with PBS and coverslipped with VectaShield fluorescent mounting medium (Vector Laboratories Inc., H-1000).

### Primary Antibodies

Primary antibodies used in this paper include rabbit polyclonal anti-human Arl13B (1:200; cat #17711-1-AP, Proteintech), rabbit polyclonal anti-human cleaved-caspase 3 (CC3, 1:250; cat #9661s, Cell Signaling Technology), mouse monoclonal anti-rat Tubulin β 3 (Tubb3, 1:500; cat #801201, BioLegend), rabbit monoclonal anti-human phospho-histone H3 (PH3, 1:200; cat #3458, Cell Signaling Technology), mouse monoclonal anti-PH3 (1:200; cat #9706, Cell Signaling Technology), and rat monoclonal anti-ZO-1 (1:50; cat #R26.4DC, DSHB).

Eight commercial antibodies purported to recognize human or mouse Togaram1 protein were tested for immunoblotting of embryonic mouse brain lysates and and immunostaining on both brain sections and dissociated mouse fibroblasts and showed either nothing or non-specific bands and staining: anti-TOGARAM1 (BiCell Scientific, cat #90704), anti-FAM179B (Invitrogen, cat #PA5-62807 and cat #PA5-65645), anti-TOGARAM1 (Atlas, cat #HPA050849), anti-FAM179B (Novus, cat #30676 and cat #58610), anti-TOGARAM1 (only tested on ICC; St. Johns Laboratory, cat # STJ195840), and anti-KIAA0423 (tested on WB only; Abnova, cat #H00023116-B01).

### Image acquisition and statistical analyses

For images in Figures 1, 2 and 3A, a Leica Microsystems MZ16F microscope with DFC300FX camera was used. A Carl Zeiss AxioObserver fluorescent widefield inverted scope microscope was used to collect images used in Figure 3D and 5C. Images from Figure 5A and 6A were collected using an inverted DeltaVision with TrueLight deconvolution microscope with softWoRx Suite 5.5 image acquisition software. A Nikon Ti2 inverted widefield scope was used to collect images from Figures 4 and 6C. ImageJ/FIJI was used for image analysis.

GraphPad PRISM software was used for statistical analyses and statistical tests of normality were performed to determine appropriate statistical test. Parametric tests were used for sample sets with normal distribution and non-parametric tests were used for sample sets without normal distribution. Specific statistical tests and sample sizes are indicated in figure legends. All error bars on graphs represent standard error of mean (SEM). No randomization or blinding was done.

## ACKNOWLEDGEMENTS

We thank the Xiaowei Lu and Ann Sutherland labs for advice, discussion, and sharing reagents. We thank Dr. Uma Paila for help searching scRNA-seq databases. We acknowledge Hayley Dingsdale for early work on the project and we thank Selena Akay for help staining and taking images. We are grateful to Kaela S. Lettieri and Klaudia Flilipek for thoughtful insights and discussion. This work was supported by NIH Grant R01HD102492 to N.D.D. and T32GM139787 Cell & Molecular Biology Training Grant to C.Q.N.

## Notes

### Competing Interest Statement

The authors have declared no competing interest.

## REFERENCES

Adams, D. J., Barlas, B., McIntyre, R. E., Salguero, I., van der Weyden, L., Barros, A., Vicente, J. R., Karimpour, N., Haider, A., Ranzani, M., Turner, G., Thompson, N. A., Harle, V., Olvera-León, R., Robles-Espinoza, C. D., Speak, A. O., Geisler, N., Weninger, W. J., Geyer, S. H., … Balmus, G. (2024). Genetic determinants of micronucleus formation in vivo. Nature, 627(8002), 130–136. 10.1038/s41586-023-07009-0

Cai, E., Barba, M. G., & Ge, X. (2023). Hedgehog signaling in cortical development. Cells, 13(1). 10.3390/cells13010021

Chee, J. M., Lanoue, L., Clary, D., Higgins, K., Bower, L., Flenniken, A., Guo, R., Adams, D. J., Bosch, F., Braun, R. E., Brown, S. D. M., Chin, H.-J. G., Dickinson, M. E., Hsu, C.-W., Dobbie, M., Gao, X., Galande, S., Grobler, A., Heaney, J. D., … Moshiri, A. (2023). Genome-wide screening reveals the genetic basis of mammalian embryonic eye development. BMC Biology, 21(1), 22. 10.1186/s12915-022-01475-0

Das, A., Dickinson, D. J., Wood, C. C., Goldstein, B., & Slep, K. C. (2015). Crescerin uses a TOG domain array to regulate microtubules in the primary cilium. Molecular Biology of the Cell, 26(23), 4248–4264. 10.1091/mbc.E15-08-0603

Di Bella, D. J., Habibi, E., Stickels, R. R., Scalia, G., Brown, J., Yadollahpour, P., Yang, S. M., Abbate, C., Biancalani, T., Macosko, E. Z., Chen, F., Regev, A., & Arlotta, P. (2021). Molecular logic of cellular diversification in the mouse cerebral cortex. Nature, 595(7868), 554–559. 10.1038/s41586-021-03670-5

Kumandas, S., Akcakus, M., Coskun, A., & Gumus, H. (2004). Joubert syndrome: review and report of seven new cases. European Journal of Neurology, 11(8), 505–510. 10.1111/j.1468-1331.2004.00819.x

La Manno, G., Siletti, K., Furlan, A., Gyllborg, D., Vinsland, E., Mossi Albiach, A., Mattsson Langseth, C., Khven, I., Lederer, A. R., Dratva, L. M., Johnsson, A., Nilsson, M., Lönnerberg, P., & Linnarsson, S. (2021). Molecular architecture of the developing mouse brain. Nature, 596(7870), 92–96. 10.1038/s41586-021-03775-x

Latour, B. L., Van De Weghe, J. C., Rusterholz, T. D. S., Letteboer, S. J. F., Gomez, A., Shaheen, R., Gesemann, M., Karamzade, A., Asadollahi, M., Barroso-Gil, M., Chitre, M., Grout, M. E., van Reeuwijk, J., van Beersum, S. E. C., Miller, C. V., Dempsey, J. C., Morsy, H., Bamshad, M. J., Nickerson, D. A., … Doherty, D. (2020). Dysfunction of the ciliary ARMC9/TOGARAM1 protein module causes Joubert syndrome. The Journal of Clinical Investigation.

Louka, P., Vasudevan, K. K., Guha, M., Joachimiak, E., Wloga, D., Tomasi, R. F.-X., Baroud, C. N., Dupuis-Williams, P., Galati, D. F., Pearson, C. G., Rice, L. M., Moresco, J. J., Yates, J. R., Jiang, Y.-Y., Lechtreck, K., Dentler, W., & Gaertig, J. (2018). Proteins that control the geometry of microtubules at the ends of cilia. The Journal of Cell Biology, 217(12), 4298– 4313. 10.1083/jcb.201804141

McNeely, K. C., & Dwyer, N. D. (2021). Cytokinetic abscission regulation in neural stem cells and tissue development. Current Stem Cell Reports, 7(4), 161–173. 10.1007/s40778-021-00193-7

Morbidoni, V., Agolini, E., Slep, K. C., Pannone, L., Zuccarello, D., Cassina, M., Grosso, E., Gai, G., Salviati, L., Dallapiccola, B., Novelli, A., Martinelli, S., & Trevisson, E. (2021). Biallelic mutations in the TOGARAM1 gene cause a novel primary ciliopathy. Journal of Medical Genetics, 58(8), 526–533. 10.1136/jmedgenet-2020-106833

Perlaza, K., Mirvis, M., Ishikawa, H., & Marshall, W. (2022). The short flagella 1 (SHF1) gene in Chlamydomonas encodes a Crescerin TOG-domain protein required for late stages of flagellar growth. Molecular Biology of the Cell, 33(2), ar12. 10.1091/mbc.E21-09-0472

Saunders, H. A. J., van den Berg, C. M., Hoogebeen, R. A., Schweizer, D., Stecker, K. E., Roepman, R., Howes, S. C., & Akhmanova, A. (2025). A network of interacting ciliary tip proteins with opposing activities imparts slow and processive microtubule growth. Nature Structural & Molecular Biology, 32(6), 979–994. 10.1038/s41594-025-01483-y

Suciu, S. K., & Caspary, T. (2021). Cilia, neural development and disease. Seminars in Cell & Developmental Biology, 110, 34–42. 10.1016/j.semcdb.2020.07.014

Thomsen, O. K., Fialová, J. L., Doganli, C., Herrera-Cid, C., Møllgård, K., Benmerah, A., Larsen, L. A., & Christensen, S. T. (2025). Primary cilia as architects of the neocortex: Roles in brain development, function, and microcephaly. Developmental Cell, 60(24), 3364–3386. 10.1016/j.devcel.2025.11.002

Wang, Y., Kraemer, N., Schneider, J., Ninnemann, O., Weng, K., Hildebrand, M., Reid, J., Li, N., Hu, H., Mani, S., & Kaindl, A. M. (2025). Togaram1 is expressed in the neural tube and its absence causes neural tube closure defects. HGG Advances, 6(1), 100363. 10.1016/j.xhgg.2024.100363

